# Characterization and remediation of sample index swaps by non-redundant dual indexing on massively parallel sequencing platforms

**DOI:** 10.1101/200790

**Authors:** Maura Costello, Mark Fleharty, Justin Abreu, Yossi Farjoun, Steven Ferriera, Laurie Holmes, Tom Howd, Tamara Mason, Gina Vicente, Michael Dasilva, Wendy Brodeur, Timothy DeSmet, Sheila Dodge, Niall J. Lennon, Stacey Gabriel

## Abstract

Here, we present an in-depth characterization of the index swapping mechanism on Illumina instruments that employ the ExAmp chemistry for cluster generation (HiSeqX, HiSeq4000, and NovaSeq). We discuss best practices for eliminating the effects of index swapping on data integrity by utilizing unique dual indexing for complete filtering of index swapped reads. We calculate mean swap rates across multiple sample preparation methods and sequencer models, demonstrating that different methods can have vastly different swap rates, and show that even non-ExAmp chemistry instruments display trace levels of index swapping. Finally, using computational methods we provide a greater insight into the mechanism of index swapping.

## INTRODUCTION

“As sequencing costs decline…” is a common phrase in the field of genomics as rapid advances in massively parallel sequencing platforms are making population scale sequencing a reality. In 2015, Illumina introduced the HiSeqX sequencer, utilizing their newest patterned flow cell and ExAmp chemistry technologies [1,2]. These instruments were purpose-built human whole genome machines, capable of producing ~1000 Gb of data per flow cell and cutting sequencing costs by two thirds over previous models. Soon after, Illumina released the HiSeq 4000 & 3000 instruments, utilizing that same patterned flow cell technology but allowing for sequencing of a wider variety of library preparation types. NovaSeq, their newest sequencer released in mid 2017, utilizes this same ExAmp chemistry with patterned flow cells but promises even higher yields of up to 3 Tb of data per flow cell by 2018 [3].

Sequencing biological samples at scale requires a highly streamlined workflow that makes the most efficient use of instrument yield and eliminates effects of lane to lane variability. To achieve the maximum cost efficiency, sample multiplexing on sequencer has become a necessity even for whole genomes. However, as others have recently reported [4-8], this new ExAmp chemistry can lead to data integrity issues due to the phenomenon of index switching or swapping. Illumina and others have reported that this swapping is likely due to residual excess free primer or adapters in the samples that, when pooled and mixed with the ExAmp reagents, can lead to spurious extension of library fragments with an oligo containing the wrong sample index. When single or combinatorial dual index schemes are used, these swapped indexes lead to read misassignment and can manifest as cross-contamination within a pool [4,5].

As this phenomenon is a property of the flow cell chemistry itself and will occur to some degree in every HiSeqX, HiSeq 4000, or NovaSeq sequencing run, the only options to completely eliminate the effects of index swapping on data integrity are to sequence one sample per lane or to use a non-redundant dual indexing strategy. As running one sample per lane is not financially feasible at scale, we needed a suitable dual index scheme. However, as of mid 2017, most vendors including Illumina still did not have a full plate, non-redundant dual indexing solution available. To create the ability to pool entire 96 well plates of dual indexed libraries, we have designed and implemented a unique dual indexing method utilizing a set of non-redundant indexes (comprising 96 unique i7 and 96 unique i5) that we have validated across multiple sample preparation workflows and Illumina sequencer models, including the NovaSeq.

## RESULTS

### Index swapping of PCR-free genomes on HiSeqX

We began multiplexing our PCR-free human whole genome libraries prior to sequencing on HiSeqX in 2015, starting with pools of 8 in February and then eventually pools of 24 by November. In our workflow, data from multiple lanes for each library are aggregated together after sequencing to achieve the desired 30X human genome coverage, and downstream analysis is performed on the aggregated data file. We chose to pool prior to sequencing for multiple reasons: (1) ease of workflow; (2) improved consistency of lane loading; and (3) reduction in the effects of lane to lane performance variability we had observed with HiSeqX flow cells. Following implementation of pooling, we began to get reports from our data users that they were observing elevated rates of sample contamination in aggregated PCR-free genome data. We ran all aggregated sample BAM (Binary Alignment/Map) files generated during the past year on HiSeqX through “VerifyBamID”, a tool designed to estimate sample % contamination in human sequencing data [9] and confirmed widespread sample contamination in PCR-free libraries at an average of 1.2%. This contamination appeared to be present in all samples generated from pooled sequencing, from all tissue sources, projects and collaborators, and appeared to worsen as we switched from 8 plex to 24 plex pools (Figure 1a).

**Figure 1:**
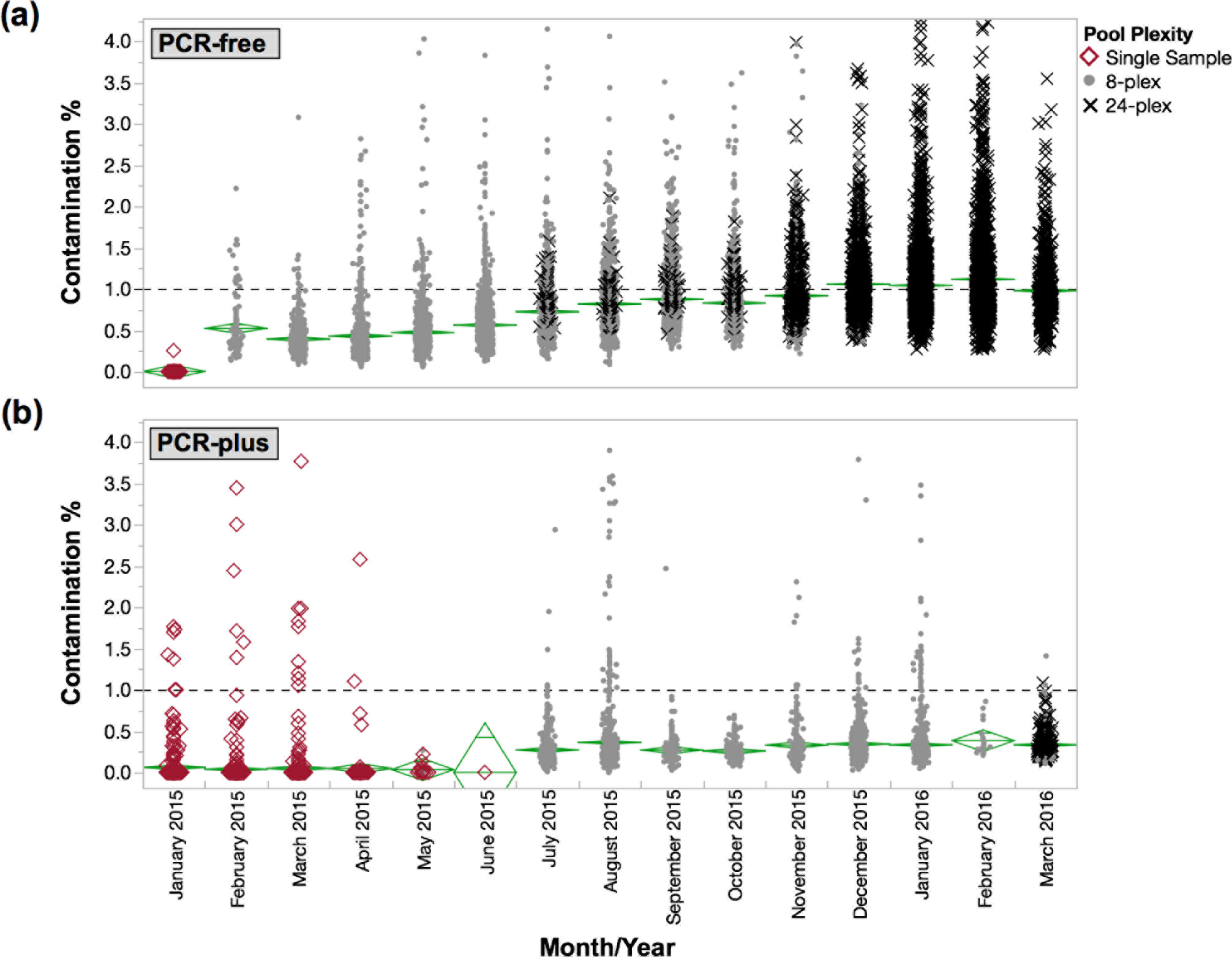
Percent contamination over time for whole genomes sequenced on HiSeqX. (a) Single indexed PCR-free library contamination by month. Contamination significantly increased when we began 8-plex pooling and worsened as we introduced 24-plex pooling. (b) Single indexed PCR-plus library contamination by month. Although overall contamination was lower for PCR-plus, rates did increase significantly as well when we began pooling.

At the same time, we were also processing a smaller number of PCR-plus whole genomes on the HiSeqX, and while we did not observe contamination at the same magnitude as PCR-free, contamination in PCR-Plus libraries had also increased significantly during the same time period (Figure 1b). Our PCR-free and PCR-plus genome libraries are made using identical protocols in the lab with the same reagents and adapter plates, by the same team, and on the same pieces of automation; the only difference is the addition of 8 cycles of PCR at the end of the process for PCR-plus. The difference in contamination rates between these two workflows provided some evidence that the contamination was likely not coming from the library preparation processes. We then took two different 24-plex PCR-free genome pools with mean library contamination rates from HiSeqX of 6.40% and 1.65% respectively, and re-sequenced these pools on MiSeq where we observed the rates of contamination for both pools dropping to 0.60% and 0.09%, respectively. This 10-fold discrepancy in contamination rate between the HiSeqX and MiSeq for the same pool of libraries strongly indicated that the contamination event was happening during the ExAmp patterned flow cell preparation or sequencing process. This observation has subsequently been reported by others [4-8].

### Non-redundant dual indexing enables identification & filtering of index swapped reads

When first released, the HiSeqX was only configured to read a single i7 library index [2]. In order to characterize what was causing this contamination, we enabled dual indexing on the HiSeqX by altering the sequencing recipes and supplementing the required i5 dual index sequencing primer. Our genome adapters were designed to be dual indexing enabled, therefore we were able to sequence a set of four 2-plex library pools containing unique combinations of dual indexes on the altered HiSeqX reading both the i7 and i5 indexes. We then ran demultiplexing and analysis on the same set of read groups from the same flow cell two different ways: first, using just the i7 single index data and second, using both the i7 and i5 dual index reads. In the data demultiplexed using just the i7 index, the contamination averaged 0.89%; however, in the data demultiplexed using both i7 and i5 indexes, the mean contamination rate dropped to just 0.13% (Supplemental Figure 2). Examining the non-demultiplexed reads from the dual indexing analysis, we observed an unusually high number of high quality reads (Q30 or greater) where the i7 and i5 indexes weren’t in the expected combinations from our standard adapter set. However, within this population of index mismatched index reads, we observed only indexes originating from libraries within the pool itself and no significant sources of outside index contamination, further indicating this wasn’t a random contamination event happening during the lab processes but something constrained to within the pool itself.

We concluded that indexes from library fragments from one sample were replacing other samples’ indexes within pools during the HiSeqX ExAmp process, which has also been confirmed by other teams [4-8]. We further hypothesize that contamination rates for single index libraries escalated as the number of samples in the pool increased because increasing the number of different libraries in an ExAmp reaction also increases the likelihood that a given library fragment will swap with a different sample’s library fragment rather than self-swapping. Unlike some [10], we have subsequently observed indexing swapping in all runs sequenced on ExAmp patterned flow cell instruments including the HiSeqX, HiSeq 4000, and NovaSeq, which agrees with Illumina’s own documentation that any sequencer model that relies on ExAmp and patterned flow cells can and will swap indexes [5]. While efficient clean up steps to remove residual index oligos may help reduce the rate of indexing swapping and computational methods may be able to compensate to some degree [11], utilizing unique non-redundant dual indexing is the only way to truly filter out swapped reads from pools sequenced on patterned flow cell data [5].

We have created and refined a novel set of 8 base pair i7 and i5 indexes, which enables a full set of 96 non-redundant dual index combinations (96 i7 indexes matched to 96 i5 indexes). This set has been validated in various sample preparation methods, and performs robustly both when used in adapter oligos ligated during library construction and in PCR primer formats that allow addition of dual indexes in targeted PCR and Nextera based protocols. They have also been screened across all Illumina sequencing platforms including random clustering and ExAmp, and 2-color or 3-color imaging, including the NovaSeq. As of September 2017, we have used this set of dual indexes on over 150,000 whole exomes on MiSeq, HiSeq 2500, and HiSeq 4000, and over 57,000 genomes on dual index enabled HiSeqX instruments, with average contamination rates close to zero (Figure 2).

**Figure 2:**
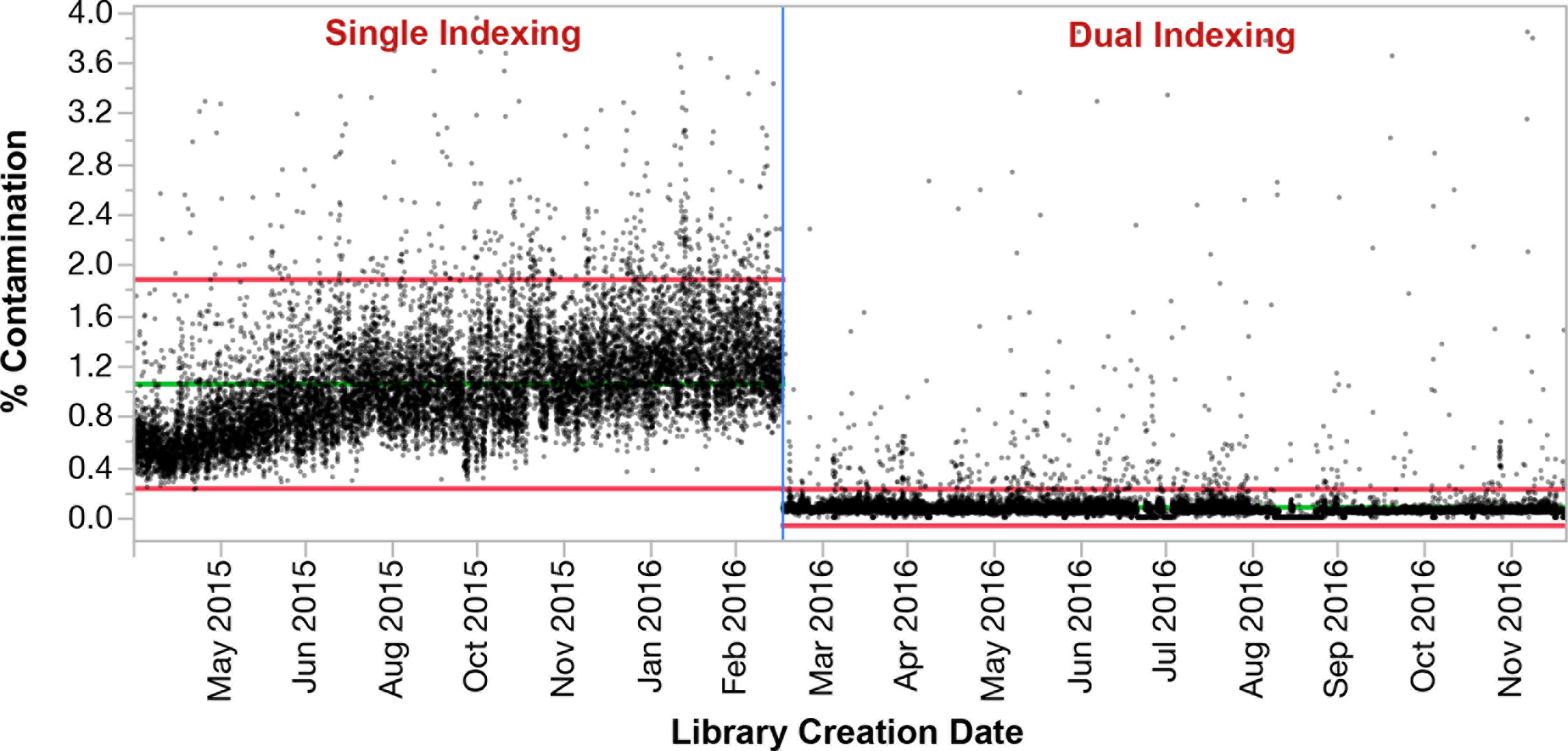
Contamination rates for single versus dual indexed pooled PCR-free libraries on HiSeqX. Percent contamination run chart by month as measured by VerifyBamID [3] for 24-plexed PCR-free genomes, demonstrating the drop in mean contamination after implementation of unique dual indexing.

### Measuring swap rates across various sample preparation methods and sequencers

As we had observed a clear difference between PCR-plus and PCR-free genome contamination rates on HiSeqX (due to swaps), we wished to survey the swap rates across our major library construction methods and across both random cluster amplification based sequencers and ExAmp patterned flow cell sequencers.

The results in Table 1 demonstrate that different library preparation methods can have drastically different swap rates and that random cluster amplification sequencers like MiSeq, NextSeq, or HiSeq 2500 can also introduce a baseline level of index swapping, although the rate is generally low [12]. While others have reported upwards of a 10% rate of index swapping on patterned flow cell sequencers [4], the highest rate of swapping we have observed is ~6% in our PCR-free workflow. Swapping at that magnitude is rare and typically we have averaged ~3% in PCR-free and ~0.25% in PCR-plus genomes. We hypothesize that PCR-free whole genomes have the highest rate of swapping due to the low yield of this particular library type following library construction. While PCR amplified libraries typically need to be diluted between 20-fold to 200-fold prior to sequencing depending on post-PCR yield, PCR-free libraries are diluted very little if at all as typical yields from that method are only a few nM of adapted library. This lack of dilution likely leads to a higher ratio of free adapter to library fragments in PCR-free pools, increasing the chance that free adapter molecules will interact with library fragments during ExAmp chemistry.

**Table 1:**
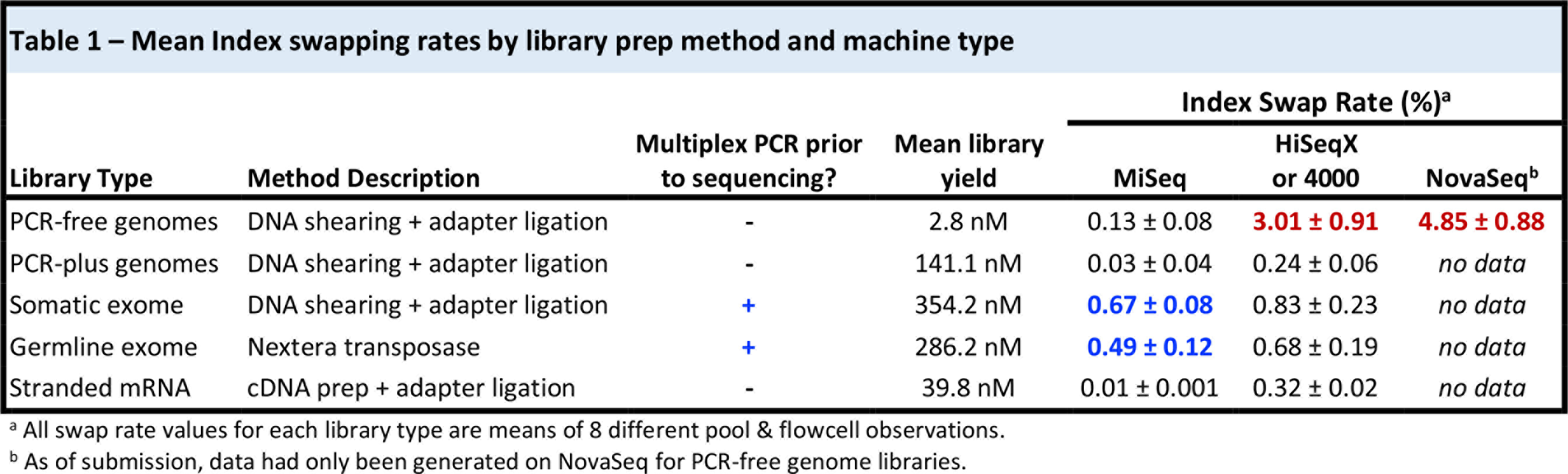
Mean Index swapping rates by library prep method and machine type.

| Library Type | Method Description | Multiplex PCR prior to sequencing? | Mean library yield | Index Swap Rate (%)<sup>a</sup> |  |  |
| --- | --- | --- | --- | --- | --- | --- |
|  |  |  |  | MiSeq | HiSeqX or 4000 | NovaSeq<sup>b</sup> |
| PCR-free genomes | DNA shearing + adapter ligation | - | 2.8 nM | 0.13 ± 0.08 | **3.01 ± 0.91** | **4.85 ± 0.88** |
| PCR-plus genomes | DNA shearing + adapter ligation | - | 141.1 nM | 0.03 ± 0.04 | 0.24 ± 0.06 | *no data* |
| Somatic exome | DNA shearing + adapter ligation | + | 354.2 nM | **0.67 ± 0.08** | 0.83 ± 0.23 | *no data* |
| Germline exome | Nextera transposase | + | 286.2 nM | **0.49 ± 0.12** | 0.68 ± 0.19 | *no data* |
| Stranded mRNA | cDNA prep + adapter ligation | - | 39.8 nM | 0.01 ± 0.001 | 0.32 ± 0.02 | *no data* |

<sup>a</sup> All swap rate values for each library type are means of 8 different pool & flowcell observations.
<sup>b</sup> As of submission, data had only been generated on NovaSeq for PCR-free genome libraries.

Interestingly, the swap rates for both germline and somatic exomes are elevated even when pools are sequenced on MiSeq. This is due to index swapping occurring during the exome capture process itself. We pool up to 12 libraries per exome hybridization reaction. Following streptavidin bead immobilization, the captured DNA is amplified off the beads in a PCR reaction. It is in this multiplex PCR where index swapping is occurring, and we have observed this previously in exome capture methods from both Agilent (Supplemental Figure 2, previously unpublished data) and Illumina (Table 1), even when sequenced on MiSeq or other random clustering sequencers.

Additionally, the standard deviation in swap rate data in Table 1 demonstrates that there can be variability of swap rates within a given library preparation method. To determine the source of this variation, we collected swap rate data on 24-plex PCR-free library pools that were sequenced across more than one flow cell, allowing us to measure variation in swap rate within an individual flow cell and across multiple flow cells containing the same pool. Index swap rates by pool and flow cell/lane are plotted in Figure 3. Swap rates for the same pool of libraries typically stay consistent across lanes within a flow cell with occasional outlier lanes, but can vary greatly between different flow cells. This data indicate that the rate of index swapping for a given pool of libraries is not entirely driven by the amount of free adapter contained in that library pool, but rather can also be influenced by ExAmp reaction setup and/or reagents and flow cell lots.

**Figure 3:**
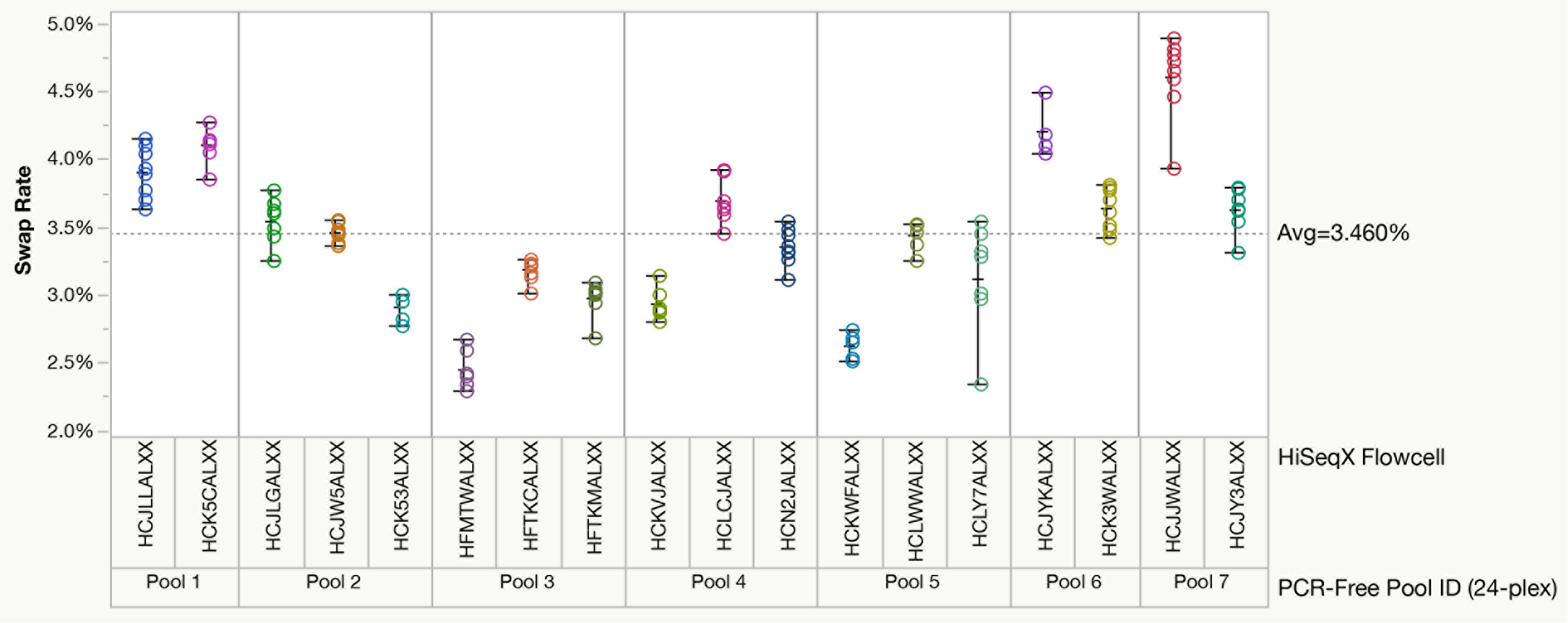
Variability of index swap rates from pool to pool and flow cell to flow cell. Index swapping rates plotted for seven 24-plex pools, each sequenced on at least two HiSeqX flow cells and prepared using identical automated methods on a Hamilton MiniStar. Each data point represents a flow cell lane. The data shows variability between different pools, but also variability for the same pools run on different flow cells, indicating that flow cell and/or ExAmp reagents also influence swap rate variability.

### Characterizing the index swapping mechanism

Utilizing unique dual indexing, we could now compare in depth swapped and unswapped reads with the goal of better understanding the mechanisms and kinetics of this phenomenon. First, we looked at a pool of 6 PCR-free dual indexed libraries sequenced on a lane of HiSeqX to determine: (i) if the rate of index swapping was the same across the entire flow cell from inlet to outlet, and (ii) if swapping occurred in a relatively normal distribution for each index or if there were biases for which indexes may be more likely to swap. The data showed that the rate of swapping was fairly uniform across both flow cell surfaces regardless of tile location (Figure 4a) and that we observed all possible index swap combinations at a relatively uniform distribution around the mean, indicating swap rates are not likely driven by amplification biases for or against certain index barcodes sequences (Figure 4b).

**Figure 4:**
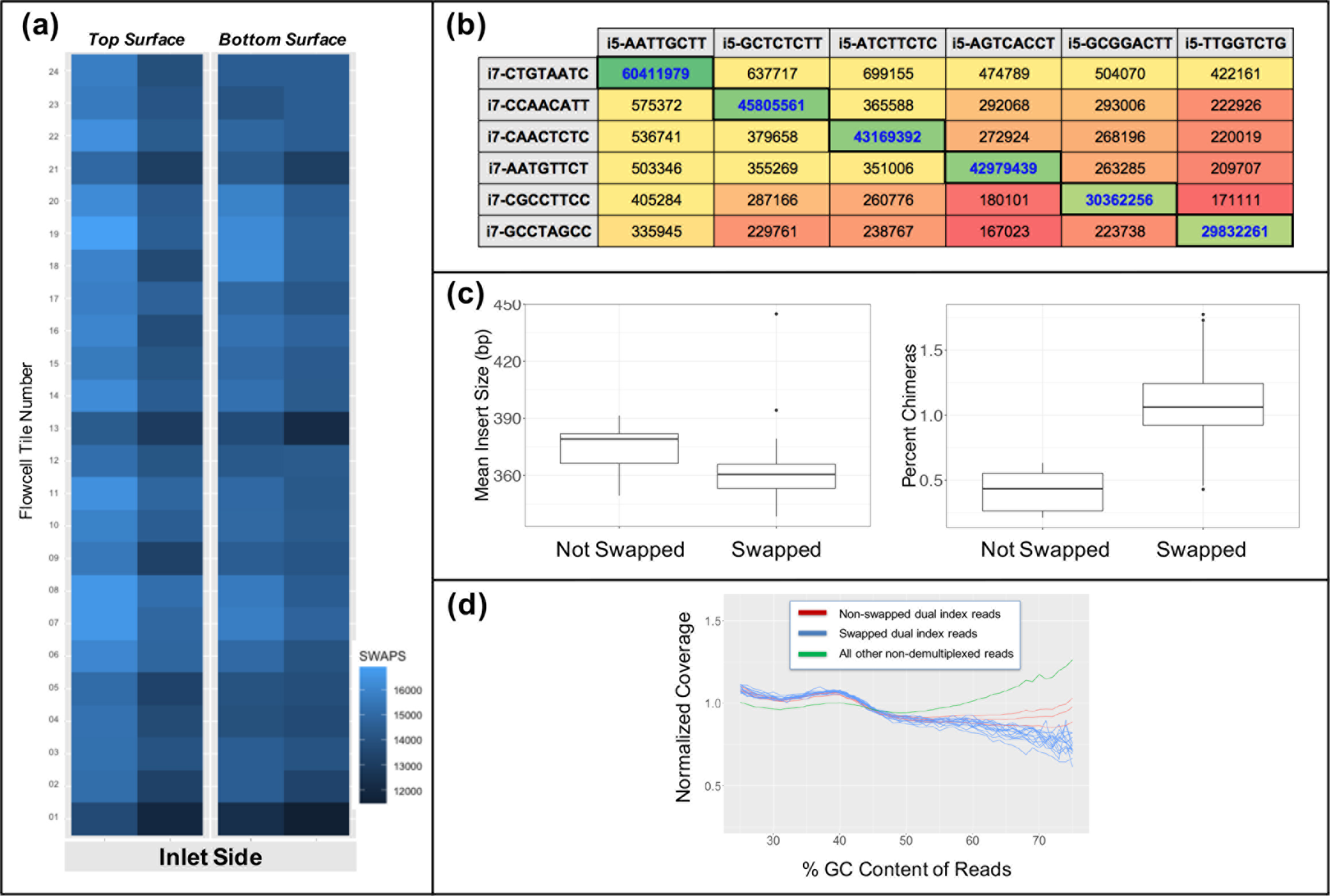
Characterization of index swapping mechanism. (a) Diagram of a HiSeqX flow cell lane colored by number of index swaps detected at each surface tile, showing relatively uniform distribution of swapping across the entire lane and both surfaces, (b) Read counts for all 36 index combinations in a 6-plex pool of uniquely dual indexed libraries. The combinations in heavy bordered cells with blue text along the diagonal are the correct index combinations; read counts for all other combinations are due to index swapping. Note all indexes participate in swapping relatively equally. (c) Mean insert size (bp) and percent chimerism calculated by Picard for both swapped and non-swapped reads. Swapped reads have shorter inserts and higher rates of chimeric read pairs. (d) Normalized human coverage across GC content bins, indicating there are less high GC reads in the swapped population (blue) compared to non-swapped (red) and all other non-demultiplexed (green) populations.

Next, we compared a variety of other sequencing and library metrics in both swapped and unswapped reads to determine if any correlations exist. We observed that swapped reads have smaller insert lengths and higher rates of chimerism than non-swapped reads (Figure 4c), as well as tend to skew towards lower % GC (Figure 4d). These observations fit the hypothesis that swapping occurs during the ExAmp chemistry step, as shorter fragments with lower %GC are known to amplify more efficiently in polymerase based amplification assays [13]. The increased rate of chimerism for swapped reads is of note for those wishing to perform structural variation detection, as these artifactual chimeric reads could be mistaken for reads derived from actual chromosomal rearrangements.

Finally, we pooled two libraries together each with unique dual indexes, one made from human DNA and one from E. coli, which allowed us to more accurately measure the rates of swapping for each of the P7/i7 and P5/i5 ends of the library fragments (Table 2). We observed that the i5 index was twice as likely to be swapped than the i7 index; however, the reason why one end of the adapter construct would preferentially be swapped over the other is unclear. Although quite rare at a rate of 0.01%, we also discovered a number of “double swaps” where both the i7 and i5 indexes from the E. coli libraries were found on human library fragments or vice versa. These double swap reads will not be removed during demultiplexing if they contain an expected combination of indexes, albeit from the wrong sample. In a non-experimental setup, this would manifest as a low rate of sample contamination.

**Table 2:**
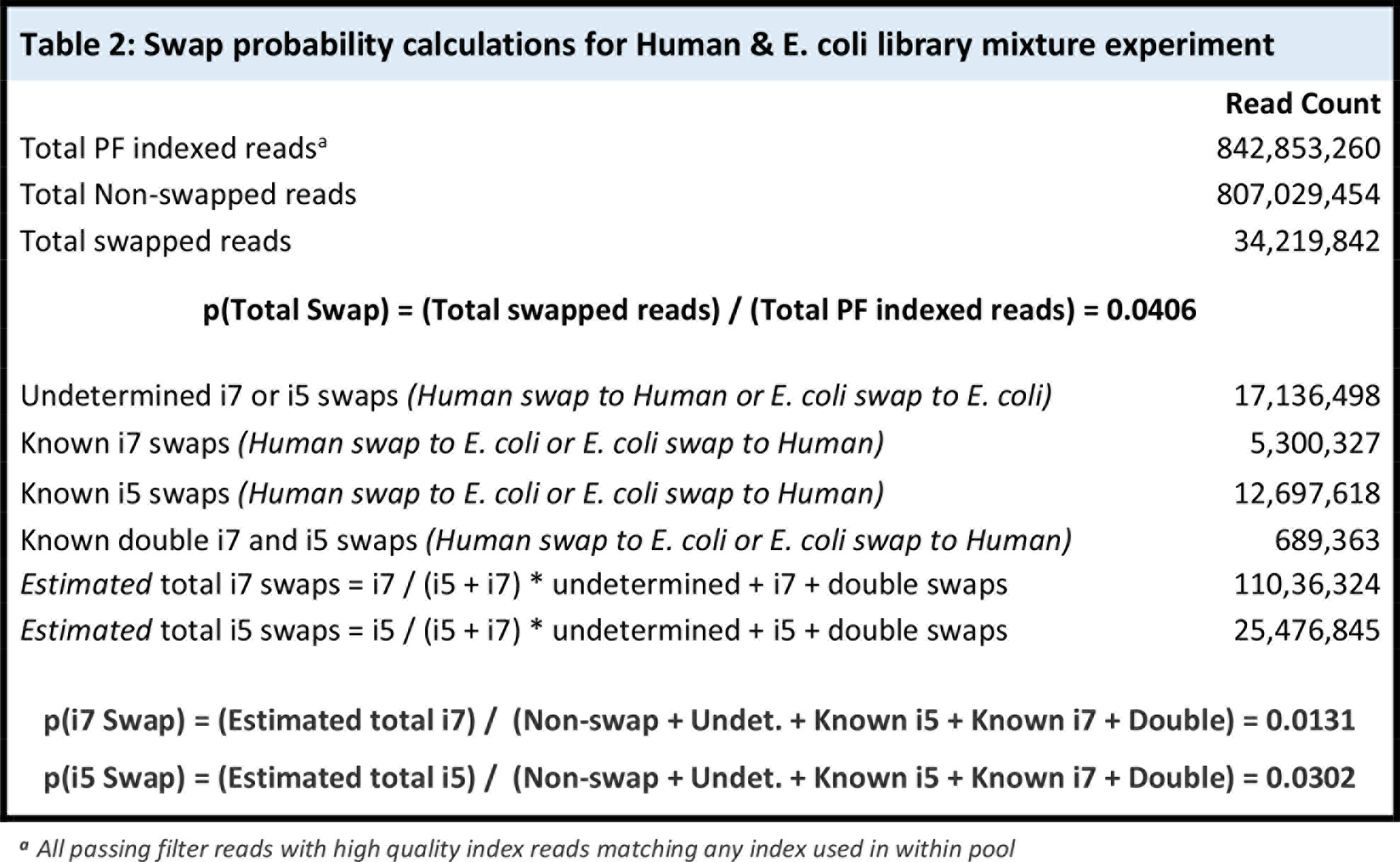
Swap Probability Calculations For Human & E. Coli Library Mixture Experiment.

Taken together, these observations indicate that index swapping is occurring relatively uniformly across all libraries within a given pool, that smaller and higher AT content fragments are more likely to be swapped as these are more efficiently amplified by the polymerase, and that this swapping phenomenon has a preference to swap the i5 index side twice as often as the i7 side.

## DISCUSSION

Our findings agree with those reported by others indicating that residual free indexing primer or adapter oligonucleotides carried over into multiplexed library pools can hybridize and extend during the ExAmp clustering chemistry leading to library fragments swapping indexes. This swapping leads to improper demultiplexing, with reads being assigned to the wrong samples which manifests as downstream read contamination in the data. We observe a minimum rate of 0.2% swapping in ExAmp/patterned flow cells across all library prep types examined. We propose the only way to effectively eliminate swapped reads from pooled sequencing data is to utilize a non-redundant dual indexing scheme and filter unexpected combinations. We believe that the elimination of index swapping is of paramount importance for those performing any sequencing studies.

It should be noted that the phenomenon of index swapping is not restricted to ExAmp chemistry from Illumina, but can occur in any scenario where multiplexed libraries are amplified together in the same vessel and residual adapters and active polymerases are present. In 2013, we had previously observed a similar phenomenon when we began pooling single indexed libraries prior to exome capture (Supplemental Figure 2) (previously unpublished data). We observed contamination levels spike, and the rate of this contamination was dependent on what PCR enzyme was used for the pooled capture PCR prior to sequencing. By implementing an early version of unique non-redundant dual indexing, we were able to filter reads with swapped indexes. Today, when we observe the rates of index swapping for exomes, we see higher than expected rates of swapping in exomes even when sequenced on random cluster amplification instruments like the MiSeq (Table 1). When designing sequencing experiments, it is therefore important to keep in mind that that any time samples are amplified together in a pool, whether in a tube during library prep or on a flow cell during ExAmp, there is a danger of index swapping induced cross contamination and non-redundant unique dual indexing should be utilized if possible.

In cases where implementation of dual indexing may be difficult or impossible for a given method, the rate of cross contamination due to index swapping contamination can be measured empirically. We recommend pooling single index libraries made from different organisms if possible, and calculating the rates of reads originating from one organism containing the i7 index from the other. Given that the rates of index swapping can vary so widely from method to method, it may be the case that the swap rate for a given method of interest may be acceptable for the type of analysis being performed and goals of a given project. However, if index swapping is occurring at a higher rate, this may lead to compromised results especially if data is to be used for detecting rare events at low allele fractions or most critically, in clinical settings.

The expected increased sequencing yields from NovaSeq will create a need for pooling larger numbers of samples for many applications to maximize cost efficiency. Major sample preparation reagent providers are just now beginning to offer unique dual indexed adapters and most do not offer full sets of 96, forcing our lab and others to operate in a “do it yourself” mode. There already exists a need for more than 96 samples worth of unique dual indexes for applications such single cell, microbial, or targeted sequencing approaches. Designing and screening new indexes can be a laborious and expensive endeavor for individual labs. As we expanded and optimized our set of dual indexes, we found it necessary to functionally validate each index pair in a costly multi-step approach, first screening indexes at the bench to ensure consistent performance in library preparation, and second ensuring that indexes sequenced as expected on multiple models of sequencers. In addition, performing quality control on incoming adapters or primers from oligo synthesis vendors can complicate implementation. Because we have observed quality issues including well to well cross contamination in index plates, we currently perform sequencing-based quality control of all incoming indexing plate batches from our oligo vendor; this is costly but highly recommended step. As the demand for indexing solutions grow, the genomics consumables industry must do better in ensuring that all labs have access to high quality and reliable indexing products that do not require the users to engage in high cost development and quality control testing activities on their own.

## CONCLUSIONS

Index swapping in pooled libraries sequenced on Illumina’s ExAmp patterned flow cell chemistry is a prevalent phenomenon that can vary in severity but that we observe is always present at a range of 0.2-6% in sequencing runs on HiSeqX, HiSeq 4000/3000, and NovaSeq. We observe that utilizing unique dual indexes for pooled libraries allows for the removal of swapped reads caused by both multiplex PCR and sequencing-chemistry induced swaps. This is particularly crucial in clinical sequencing settings or in single cell sequencing, where even low percentages of anomalous reads are unacceptable.

The phenomenon of index swapping was discovered and reported by the genomics technology user community, a fact that reinforces the need to be vigilant and closely review any novel sample preparation or sequencing technologies from both new and well established companies. We should challenge our technology vendors to ensure they have performed thorough validation of new technologies prior to their release, and further ensure that we have the tools at our disposal to monitor and ensure the integrity of sequencing data.

## MATERIALS AND METHODS

### Preparation of sequencing libraries

Library construction was performed using Kapa Biosystems reagents as described by Fisher et al. [14] with some slight modifications. For whole genomes, initial genomic DNA input was reduced from 3μg to 250ng for PCR-free or 50 ng for PCR-plus. For germline exomes, input to Nextera based library prep was 50 ng. For somatic exomes, DNA input into sheared library prep was 100 ng. Subsequent exome capture for both somatic and germline exomes were performed using the Illumina exome oligo pool with a 38 Mb target design. For stranded RNA-seq, 250 ng of total RNA was used as input into the TruSeq stranded mRNA sequencing kit (Illumina). Dual indexed library oligos were custom-ordered from IDT. For ligation adapters, these were ordered HPLC purified, pre-annealed, and in single use plates each at a concentration of 15 uM. For Nextera PCR primers, these were ordered standard desalted, forward and reverse premixed, and in single use plates at a concentration of 10 uM.

Following sample preparation, libraries were quantified using quantitative PCR (kit purchased from KAPA biosystems) with probes specific to adapter ends in an automated fashion on Agilent’s Bravo liquid handling platform. Based on qPCR quantification, libraries were normalized and pooled on the Hamilton MiniStar liquid handling platform. For HiSeqX and HiSeq4000, pooled samples were normalized to 2 nM and denatured with 0.1N NaOH for a loading concentration of 200 pM. For Novaseq, pooled samples were normalized 1 nM and denatured with 0.1N NaOH in for a loading concentration of 200 pM. For MiSeq, pooled samples were normalized to 2 nM and denatured with 0.1N NaOH for a loading concentration of 14 pM.

### Cluster amplification and sequencing

Cluster amplification of denatured templates and paired-end sequencing was performed according to the manufacturer’s protocol (Illumina) for HiSeq X, HiSeq 4000, NovaSeq, or MiSeq. For single index sequencing, an additional 8-bp i7 index read was sequenced. For dual index, additional 8-bp i7 index and 8-bp i5 index reads were sequenced. Dual indexed sequencing on HiSeqX was initially enabled outside of standard control software versions and kits were supplemented with dual index primer HP14. As of October 2017, Illumina is officially supporting dual indexing on HiSeqX via HiSeq control software v3.5.0 and inclusion of HP14 in revised reagent kits.

### Sequencing data analysis

Output from Illumina software was processed by the Picard data-processing pipeline to yield BAM files containing quality-calibrated, aligned reads. Contamination was calculated using VerifyBamID [9]. Index swapping calculations were made by tabulating per-tile index read information to determine the percentages of both correct and swapped dual-indexed combinations present.

For experiments comparing metrics for swapped versus non-swapped reads, 12 libraries of NA12878 human DNA and 12 libraries of E. Coli-K12_MG12655 were prepared as PCR-free genome libraries with unique dual index combinations. The Picard data-processing pipeline was then used to aggregate bam files while allowing all possible barcode combinations. Since there are 24 possible barcodes (for both i5 and i7), there are 24^2=576 possible pairs of barcodes. Each possible barcode pair was aggregated into its own bam file resulting in 576 BAM files, of which 552 constitute swaps. We then aligned these bams to a reference containing both human and E. Coli contigs and counted the number of reads mapping to human and E. coli respectively from each file. Insert size, the rate of chimerism, and GC content were then calculated independently for the swapped and non-swapped BAMs.

We assume that all reads were mapped correctly to their organism of origin, and from this we are able to determine the location where a swap occurred, i5 or i7 index. When reads and barcodes belong to the same species we can identify that a swap has occurred, however we cannot determine where the swap occurred. When barcodes originally belonging to differing organisms are found on a read, we can identify the barcode that swapped by assuming that the unique barcode belonging to the organism the read maps to did not swap. Likewise, it is possible to identify cases where both the i5 and i7 barcodes swapped when a read maps to a particular organism, but both its barcodes point to a sample from the other organism. Using this simple approach we were able to estimate the probability of swaps occurring at each end of the library fragment.

